# Non-Invasive monitoring of normal tissue radiation damage using quantitative ultrasound spectroscopy

**DOI:** 10.1101/2022.05.10.491268

**Authors:** Marjan Rafat, Ahmed El Kaffas, Ankush Swarnakar, Anastasia Shostak, Edward E. Graves

## Abstract

**Background:** While radiation therapy (RT) is an critical component of breast cancer therapy and is known to decrease overall local recurrence rates, recent studies have shown that normal tissue radiation damage may increase recurrence risk. Fibrosis is a well-known consequence of RT, but the specific sequence of molecular and mechanical changes induced by RT remains poorly understood.

**Purpose:** There is currently a need to understand the contribution of the irradiated tissue microenvironment to recurrence to improve cancer therapy outcomes. This study seeks to evaluate the use of quantitative ultrasound spectroscopy (QUS) for real time determination of the normal tissue characteristic radiation response and to correlate these results to molecular features in irradiated tissues.

**Methods:** Murine mammary fat pads (MFPs) were irradiated to 20 Gy, and QUS was used to analyze tissue physical properties pre-irradiation as well as at 1, 5, and 10 days post-irradiation. Tissues were processed for scanning electron microscopy imaging as well as histological and immunohistochemical staining to evaluate morphology and structure.

**Results:** Tissue morphological and structural changes were observed non-invasively following radiation using mid-band fit (MBF), spectral slope (SS), and spectral intercept (SI) measurements obtained from QUS. Statistically significant shifts in MBF and SI indicate structural tissue changes in real time, which matched histological observations. Radiation damage was indicated by increased adipose tissue density and extracellular matrix (ECM) deposition.

**Conclusions:** Our findings demonstrate the potential of using QUS to non-invasively evaluate normal tissue changes resulting from radiation damage. This supports further pre-clinical studies to determine how the tissue microenvironment and physical properties change in response to therapy, which may be important for improving treatment strategies.

## 1. Introduction

Radiation therapy (RT) has been shown to reduce rates of local tumor recurrence in breast cancer. However, irradiation of adjacent normal tissues continues to be a limiting factor. Normal tissue irradiation is known to induce pathological fibrosis and tissue stiffening [*1*]. Recent studies have found that irradiation of normal tissues enhances tumor cell recruitment and may correlate to breast cancer recurrence [*2, 3*]. The role of tissue mechanical and structural properties in facilitating recurrence is unknown. Tissue stiffness is often determined in terminal experiments using techniques such as atomic force microscopy (AFM) [*4*]. Methods to non-invasively monitor tissue properties and changes in the tissue microenvironment over time may lead to insights about the extent of damage, level of fibrosis, and other post-treatment conditions promoting recurrence.

Previous studies used quantitative ultrasound spectroscopy (QUS) to determine tissue abnormalities and tumor types [*5, 6*] and tumor response to therapy [*7-9*]. QUS works by parameterizing frequency-dependent backscatter. Spectral QUS parameters, including mid-band fit (MBF), spectral slope (SS), and spectral intercept (SI), have been used to detect tissue microsctructural and morphological properties, cell death, and microscopic stiffness changes [*5, 10-12*]. In this study, QUS measurements were taken at multiple timepoints following mouse mammary fat pad (MFP) irradiation to investigate the normal tissue radiation response. We found significant QUS parameter changes following RT, indicating soft tissue changes. Thus, QUS may be used pre-clinically to study how microenvironmental properties correlate to recurrence propensity in breast cancer.

## 2. Methods and Materials

### Radiation

Animal studies were performed in accordance with institutional guidelines and protocols approved by the Stanford University Institutional Animal Care and Use Committee. MFPs of 8-10 week old female BALB/c mice (n=5, Charles River Laboratories) were irradiated using a 250kVp cabinet x-ray system filtered with 0.5mm Cu. Mice were anesthetized with 80mg/kg ketamine hydrochloride and 5mg/kg xylazine i.p. and shielded using a 3.2mm lead jig with 1cm circular apertures to expose the left #4 MFP to 20 Gy. 5 corresponding control mice were anesthetized but were not irradiated. Transmission through the shield was approximately 1%.

### Ultrasound Imaging and QUS

Treated and control animals were imaged with 3D high-frequency ultrasound immediately pre-treatment and after 1, 5, and 10 days post-treatment (**Figure 1A**). Mouse fur was removed under isoflurane anesthesia using with electric clippers (Pet Trimmer Pocket Pro, Wahl, Sterling, IL). Ultrasound images were collected at 40MHz with an MS-550D (RMV-710B: ∼70µm axial resolution, ∼140µm lateral resolution, focal length of 15mm) transducer on a VEVO2100 system (Visualsonics, Toronto, ON). Mice anesthetized with isoflurane were placed on the dorsal side and covered with warm gel; the transducer was then placed over the MFP and swept over the region at a step size of 0.1mm using a motorized scan stage. The focal zone was placed in the center of the left #4 MFP.

**Figure 1.**
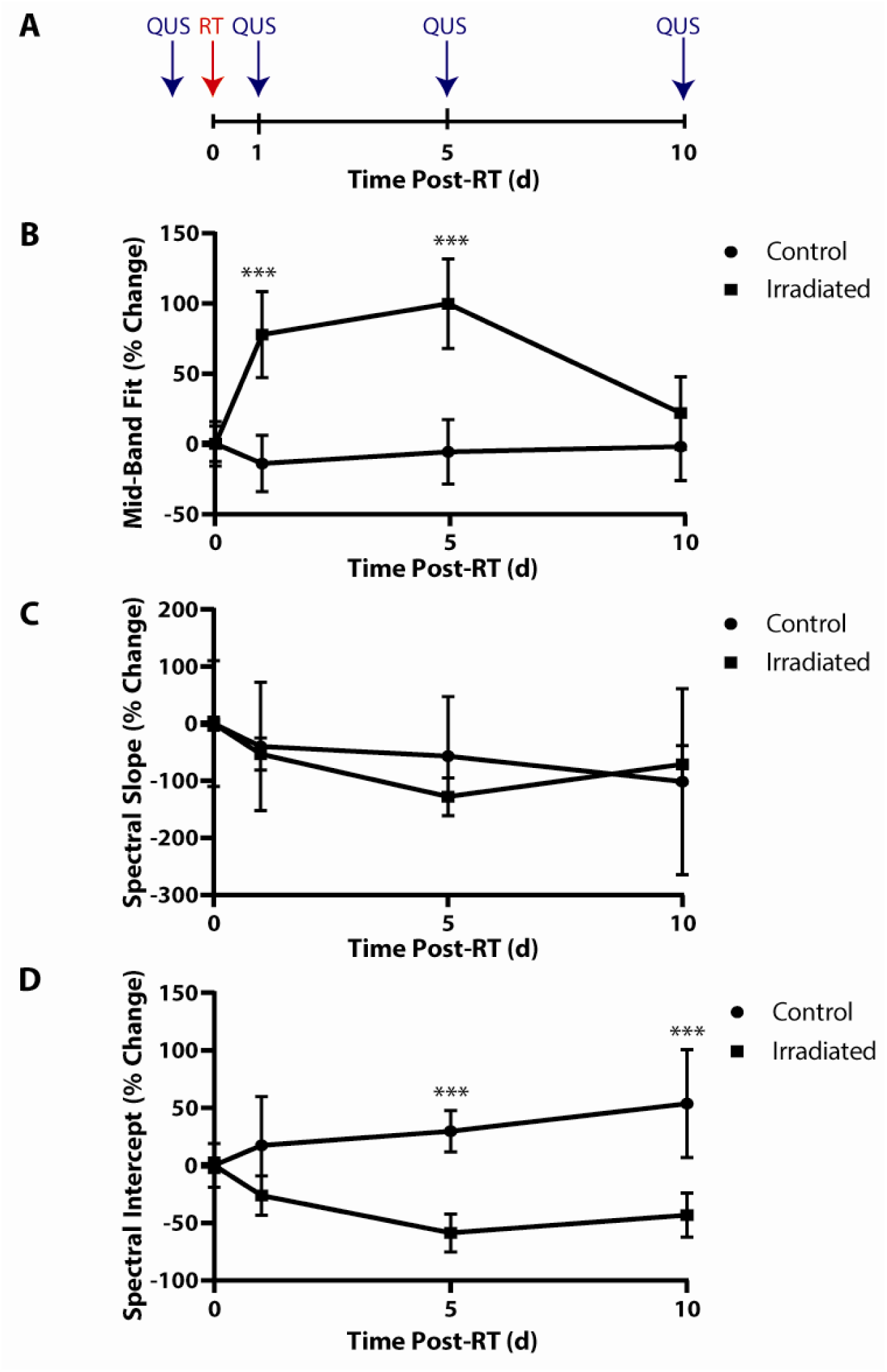
QUS measurements following irradiation of mouse MFPs reveal an increase in stiffness. (A) Experimental timeline. Mouse mammary fat pads were irradiated to 20 Gy. Ultrasound imaging was conducted immediately before irradiation and 1, 5, and 10 days post-irradiation. Histology was performed 10 days post-irradiation to correlate QUS measurements with radiation response. Percent changes in mid-band fit (B), spectral slope (C), and spectral slope (D) across 10 days for control and irradiated groups. Error bars show standard error with ***p<0.001 as determined by one-way ANOVA analysis between the control and irradiated groups.

Analysis of RF data was performed using previously validated QUS methodologies [*8, 13, 14*]. Briefly, regions-of-interest (ROIs) were selected to encompass the MFP and exclude image artifacts and skin lines. A Fourier transform was applied to the RF data along each scan line within the ROI to obtain a power spectrum; spectra were averaged over all scan lines in the ROI and normalized to a reference phantom power spectrum [*15-20*]. Linear regression analysis was performed on each normalized power spectrum to generate a best-fit-line within a -6dB window to obtain MBF, SS, and SI parameters [*15-20*]. SS and SI represent the slope and the 0-MHz intercept of the best-fit-line, respectively. The MBF is the solution of the best-fit-line for the power spectrum at the center frequency of the -6dB bandwidth window. Structural and morphological properties impact these parameters, which are related to the scatterer size (SI, SS), the acoustics scatterer concentration (MBF, SI), or difference in acoustic impedance between the scatterer and its surrounding medium (SI). Spatial parametric images of the MBF were generated by displaying the results of a sliding window analysis on a pixel-by-pixel basis within the ROI [*17, 19, 20*]. Statistical analysis was done with a multivariate model analysis and a one-way ANOVA as presented in **Tables 1 and 2**.

**Table 1.**
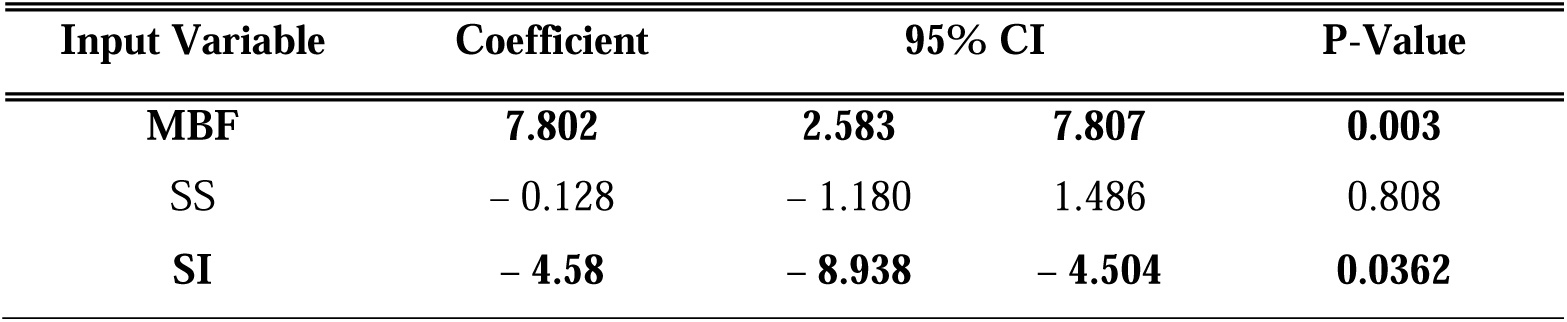
Generalized multivariate linear regression model with spectral parameters (MBF, SS, and SI) for tissue classification. The model’s regression coefficients for each variable are reported. A 95% confidence interval (CI) and P values (bolded when significant with p < 0.05) are also shown.

| Input Variable | Coefficient | 95% CI |  | P-Value |
| --- | --- | --- | --- | --- |
| MBF | **7.802** | **2.583** | **7.807** | **0.003** |
| SS | - 0.128 | - 1.180 | 1.486 | 0.808 |
| SI | - **4.58** | - **8.938** | - **4.504** | **0.0362** |

### Histological Analysis

Following ultrasound imaging 10 days post-RT, tissues were removed and placed in 10% formalin for 24hr (4° C). Fixed MFPs were placed in 70% ethanol before embedding in paraffin. Sections (4μm) were deparaffinized, rehydrated, and stained with hematoxylin and eosin (H&E) (Richard-Allan Scientific). For immunofluorescence, deparaffinized and rehydrated sections were boiled in citric acid (10mM, pH 6) for antigen retrieval and treated with 3% H_2_O_2_. Blocking in 10% goat serum was followed by incubation overnight (4° C) with an anti-perilipin-1 (1:200; Abcam) primary antibody followed by incubation with a fluorescent secondary antibody. Sudan Black B was used to quench autofluorescence before nuclear staining with 4′,6-diamidino-2-phenylindole dihydrochloride. Samples were imaged using an inverted Leica microscope (DMi8). ImageJ was used to quantify cell number and adipocyte size.

### Scanning Electron Microscopy (SEM)

SEM was performed at Stanford’s Cell Sciences Imaging Facility. MFPs from a separate group of mice (n=5 control and irradiated) were removed 10 days post-RT and placed in 2% glutaraldehyde with 4% paraformaldehyde in 0.1M sodium cacodylate buffer (pH 7.4) fixative at 4° C until placed in 1% aqueous OsO_4_ for 1hr (room temperature), washed with dI H_2_O, and dehydrated serially in ethanol (50%, 70%, 90%, 100%) before critical point drying with liquid CO_2_ (Auto Samdri 815 Series A, Tousimis, Rockville, MD) and coated with Au/Pd for 2min with 40mA with a sputter coater (Denton Desk II). MFPs were imaged using SEM at 1.5kV (Zeiss Sigma).

## 3. Results

### Ultrasound Spectral Analysis

To determine wheter QUS parameters could detect changes in tissue properties following radiation, a logistic regression was performed (**Table 1**). MBF and SI were found to be statistically significant indicators of radiation-induced tissue changes. In control MFPs, MBF did not change significantly over time. In irradiated tissues, MBF increased from day 0 to day 5 post-RT by 98% (p<0.001) (**Figure 1B**), suggesting tissue structural changes. From day 5 to 10 following irradiation, MBF decreases. SS shows a shallow decrease from day 0 to 10 in control tissues while a decreasing trend was observed up to day 5 in treated tissues (**Figure 1C**). Large error bars in the SS data indicate that changes in the control tissues may be attributed to extraneous variables, including cell growth and image artifacts. SI increased consistently from day 0 to 10 in unirradiated tissues (**Figure 1D**). Between days 0 to 5 post-RT, SI decreased in irradiated tissues. SS and SI increased from day 5 to 10 in irradiated groups (**Figures 1C, D**). Statistical significance was determined between the control and irradiated groups for each timepoint and spectral parameter using one-way ANOVA analysis.

### Adipose Tissue Morphology

H&E staining in control and irradiated tissues was performed to evaluate changes in tissue morphology 10 days post-RT, and representative images are shown in **Figure 2A**. An increase in adipocyte density due to a reduction of adipocyte size as well as an increase in extracellular matrix (ECM) deposition indicative of fibrosis following irradiation was observed. Staining of the adipocyte-specific perilipin-1 confirmed this trend (**Figure 2B**). Radiation caused a statistically significant increase in cell number (**Figure 2C**), correlating to the shifts in QUS parameters between 5-10 days post-RT, and a decrease in average adipocyte size (**Figure 2D**), indicating tissue damage.

**Figure 2.**
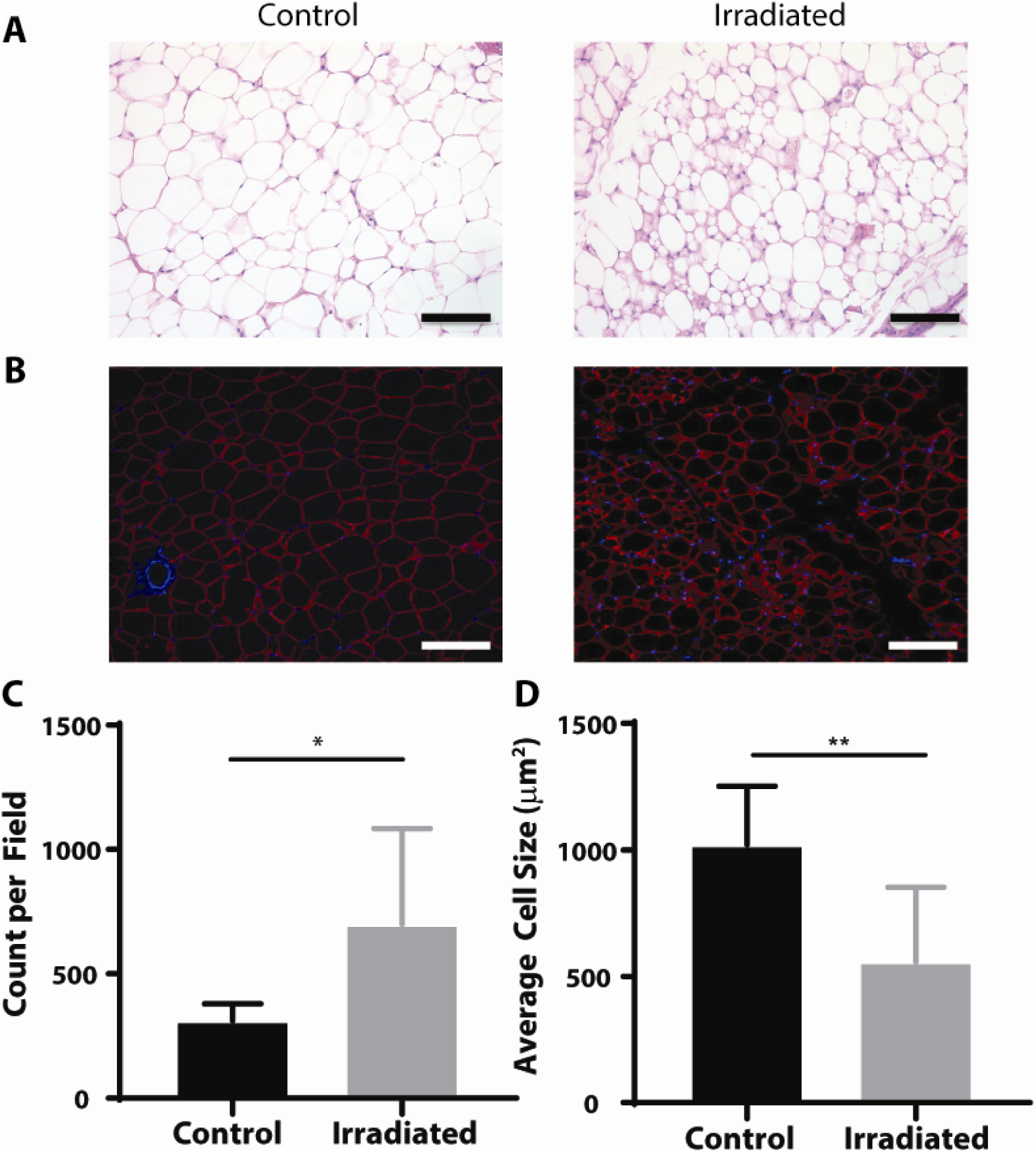
Adipocyte morphology change following irradiation. (A) Representative hematoxylin and eosin (H&E) staining of control (n=5) and irradiated MFPs (n=5). (B) Perilipin (red) and nuclear (blue) staining of control and irradiated MFPs demonstrate an increase in adipocyte density in irradiated MFPs. Quantification of (C) adipocyte count per field and (D) average adipocyte size. Error bars represent standard error with *p < 0.05 and **p < 0.01 as determined by a two-tailed unpaired t-test. Scale bar is 100 μm.

We used SEM to visualize changes in the MFP following radiation damage (**Figure 3**). We observed large, uniform adipocytes with minimal ECM deposition in control unirradiated tissues (**Figure 3A**). Radiation not only altered the uniformity of adipocyte size and shape but also enhanced ECM and collagen deposition and fiber disorder (**Figure 3B**). This suggests that irradiated MFPs became fibrotic 10 days following irradiation as expected.

**Figure 3.**
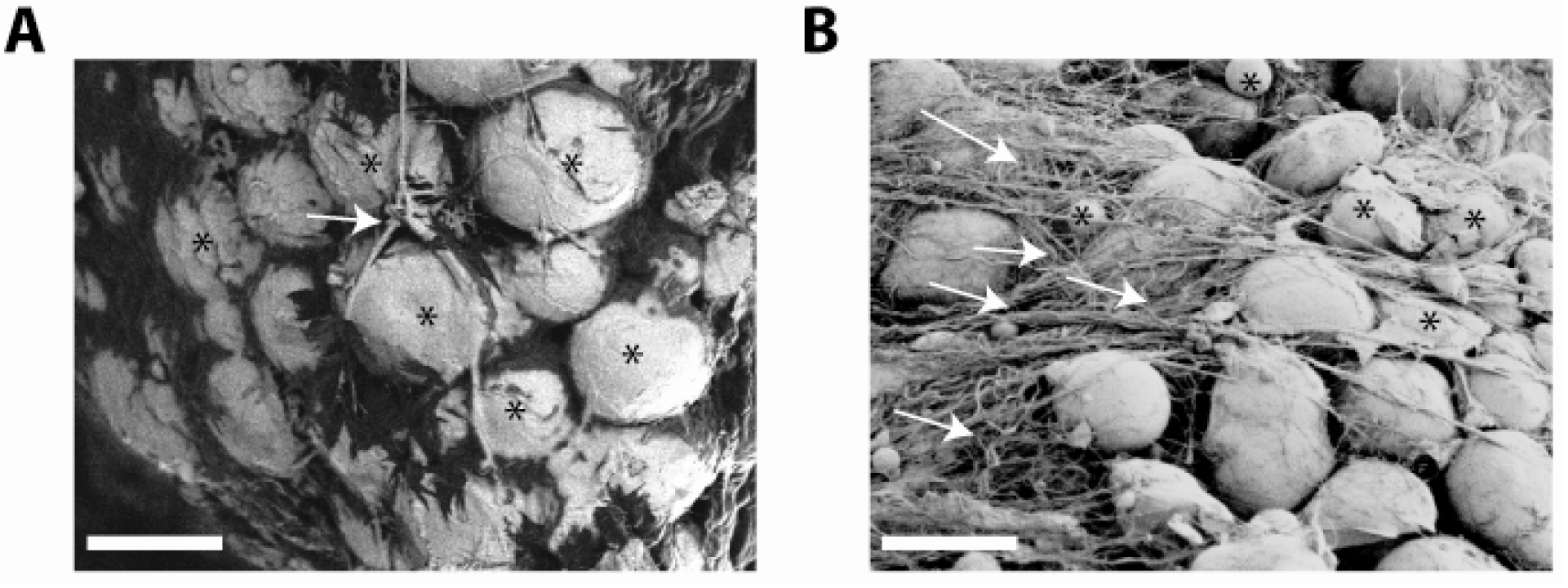
Visualizing tissue properties following irradiation of MFPs shows an increase in ECM deposition. Scanning electron microscopy was used to visualize the MFP in (A) unirradiated and (B) irradiated mouse tissues. White arrows indicate ECM deposition, and black stars denote adipocytes. An increase in ECM deposition, fiber disorder, and a decrease in adipocyte size were observed in irradiated MFPs. Scale bar is 50 μm.

## 4. Discussion

We used QUS in this proof of concept study to evaluate changes in soft tissues damaged by irradiation. We examined normal tissue physical and morphological properties serially up to 10 days after irradiation as these conditions induced tumor cell recruitment in previous studies [*3*]. MBF and SI detected tissue structural changes, potentially correlating to stiffening, and cell death up to 5 days following irradiation (**Figure 2**). A reversal in the trend for MBF, as well as a change in slope for SI between 5 and 10 days post-RT, may correlate to potential death of the irradiated cells and replacement by healthy infiltrating cells. This agrees with previous data showing macrophage infiltration into irradiated murine MFPs [*3*]. A limitation of this study is that histology and SEM were performed only at the experimental endpoint. However, QUS trends over 10 days complemented observed changes in adipocyte morphology and density as well as ECM deposition and fibrosis (**Figures 3, 4**).

Investigating tissue mechanical and structural properties due to therapy response may be critical for analyzing conditions conducive to tumor cell migration, invasion, and retention in radiation-damaged sites. Future experiments will link these dynamic QUS parameters to matched histological timepoints, expression of damage markers, and AFM measurements for model validation. Taken together, our work demonstrates that QUS enables non-invasive monitoring of normal tissue properties following RT, which can be extended to evaluating the tissue response to surgery, chemotherapy, and immunotherapy.

## 5. Conclusions

This work establishes the potential for using QUS to evaluate real-time normal tissue property changes in response to radiotherapy. We analyzed the spectral parameters of MBF, SS, and SI to determine normal tissue stiffness and cell death up to 10 days post-RT, which correlated to morphological and structural changes in the irradiated MFP. Utilizing an ultrasound-based approach to non-invasively examine tissue properties may serve as a valuable tool for determining radiation response in the tissue microenvironment.

## Acknowledgments

This research was financially supported by the Katherine McCormick Advanced Postdoctoral Fellowship, NIH grant# K99/R00CA201304, and the Ruth L. Kirschstein National Research Service Award PA-14-015 Grant# T32CA121940 (M.R.).

## Data Availability Statement

Research data are stored in an institutional repository and will be shared upon request to the corresponding author.

